# MicroRNA levels in bone and blood change during bisphosphonate and teriparatide therapy in an animal model of postmenopausal osteoporosis

**DOI:** 10.1101/591990

**Authors:** Roland Kocijan, Moritz Weigl, Susanna Skalicky, Elisabeth Geiger, James Ferguson, Gabriele Leinfellner, Patrick Heimel, Peter Pietschmann, Johannes Grillari, Heinz Redl, Matthias Hackl

**Affiliations:** Ludwig Boltzmann Institute for Experimental and Clinical Traumatology in AUVA research center; Donaueschingenstraße 13, 1200 Vienna, Austria; Hanusch Hospital, 1st Medical Department, Heinrich Collin-Str. 30, 1140 Vienna, Austria; TAmiRNA GmbH, Muthgasse 18, 1190 Vienna, Austria; Austrian Cluster for Tissue Regeneration, Austria; Karl Donath Laboratory for Hard Tissue and Biomaterial Research, Department of Oral Surgery, Medical University of Vienna, Austria; Department of Pathophysiology and Allergy Research, Center for Pathophysiology, Infectiology and Immunology, Medical University of Vienna, Vienna, Austria; Christian Doppler Laboratory on Biotechnology of Skin Aging, Department of Biotechnology, BOKU - University of Natural Resources and Life Sciences Vienna, Vienna, Austria

**Keywords:** bone microstructure, circulating microRNA, osteoporosis, bisphosphonate, parathyroid hormone (PTH), μCT, RUNX2

## Abstract

MicroRNAs control the activity of a variety of genes that are pivotal to bone metabolism. Therefore, the clinical utility of miRNAs as biomarkers and drug targets for bone diseases certainly merits further investigation. This study describes the use of an animal model of postmenopausal osteoporosis to generate a comprehensive dataset on miRNA regulation in bone tissue and peripheral blood during bone loss and specifically anti-resorptive and osteo-anabolic treatment.

Forty-two Sprague-Dawley rats were randomized to SHAM surgery (n=10) or ovariectomy (OVX, n=32). Eight weeks after surgery, OVX animals were further randomized to anti-resorptive treatment with zoledronate (n=11), osteo-anabolic treatment with teriparatide (n=11), or vehicle treatment (n=10). After 12 weeks of treatment, bone and serum samples were used for microRNA analysis using next-generation sequencing (NGS), mRNA levels using RT-qPCR, and bone microarchitecture analysis using nanoCT.

Ovariectomy resulted in loss of trabecular bone, which was fully rescued using osteo-anabolic treatment, and partially rescued using anti-resorptive treatment. NGS revealed that both, anti-resorptive and anabolic treatment had a significant impact on miRNA levels in bone tissue and serum: out of 426 detected miRNAs, 46 miRNAs were regulated by teriparatide treatment an d 10 by zoledronate treatment (p-adj. < 0.1). Interestingly, teriparatide and zoledronate treatment were able to revert miRNA changes in tissue and serum of untreated OVX animals, such as the up-regulation of miR-203a-3p, a known osteo-inhibitory miRNA. We confirmed previously established mechanisms of miR-203a by analyzing its direct target Dlx5 in femoral head.

Our data reveal a significant effect of ovariectomy-induced bone loss, as well as the two major types of anti-osteoporotic treatment on miRNA transcription in femoral head tissue. These changes are associated with altered activity of target genes relevant to bone formation, such as Dlx5. The observed effects of bone loss and treatment response on miRNA levels in bone are also reflected in the peripheral blood, suggesting the possibility of minimally-invasive monitoring of bone-derived miRNAs using liquid biopsies.

**Highlights:**

1. microRNA expression in bone tissue is altered by osteo-anabolic and anti-resorptive therapy in OVX rats.
2. microRNA changes in untreated OVX rats are reverted by anti-osteoporotic therapy.
3. miR-203a is up-regulated during bone loss and down-regulated following therapy.
4. Bone tissue and serum levels of miR-203a are highly correlated.

## 1. INTRODUCTION

Osteoporosis is a worldwide affliction associated with low-traumatic fractures, morbidity and mortality ^(1–3)^. Currently available biomarkers for the diagnosis and follow-up of bone diseases are bone turnover markers (BTMs) ^(4)^. BTMs reflect bone resorption and bone formation and therefore the activity of bone cells, and are recommended for monitoring of treatment response ^(5)^. MicroRNAs (miRNAs) are a novel group of biomarkers ^(6)^. miRNAs are small non-coding RNAs that are involved in several biological processes such as stem cell differentiation, cell cycle control, apoptosis, aging and bone metabolism ^(7–10)^. In regards to bone metabolism and bone health, several miRNAs have been identified as potent regulators of osteogenic differentiation and osteoclast formation ^(11)^. Interestingly, genetic association studies have also identified polymorphisms in miRNA binding sites in 3’UTRs of FGF2, ON, and COL1A2 to be associated with bone mineral density ^(12)^. Vice-versa, individuals carrying WNT1 mutations and thus exhibiting severe bone loss and multiple fractures exhibit particular miRNA profiles in their serum ^(13)^, including down-regulation of miR-31-5p, a putative regulator of WNT1. In addition, miR-31-5p was shown to inhibit bone formation via RUNX2 and SATB2 ^(14)^, while inhibition of miR-31-5p activity using antisense RNAs was shown to improve fracture healing in vivo ^(15)^. Finally, using ovariectomized mice, changes in expression of miRNAs and mRNAs in femur and tibia were identified already after a period of 4 weeks, and found to be associated with reduced bone formation ^(16)^.

Hence, circulating miRNAs have been proposed as potentially valuable biomarkers for the diagnosis of osteoporosis and the prediction of low-traumatic fractures ^(6)^, based on studies performed in women with postmenopausal osteoporosis with and without low-traumatic fractures ^(17–19)^, and the observation that circulating miRNAs are associated to bone microstructure and histomorphometry ^(20)^.

Although we recently observed in a cross-sectional study that previous anti-resorptive therapy did not significantly alter serum miRNA levels in osteoporotic individuals, the effect sizes observed for treated versus untreated subjects suggested a potential response of miRNA biomarkers to therapeutic intervention ^(21)^. In addition, a recent study performing targeted analysis of a small set of circulating miRNAs in patients undergoing denosumab or teriparatide treatment revealed several significant changes ^(22)^. While bisphosphonates still represent the most commonly prescribed type of osteoporosis drug, there are no data describing drug effects on miRNA levels in bone tissue and circulation. Nevertheless, such data are important to validate the clinical utility of novel bone biomarkers for diagnosis and monitoring response to treatment in bone diseases and could potentially unravel novel drug targets for intervention. Accordingly, we designed an animal trial that would enable us to systematically analyze miRNA levels in bone tissue and serum during bone loss and therapeutic intervention. We specifically considered the analysis of common changes in bone-related miRNAs at tissue and cell-free blood level (serum) after a 12-week treatment course, as well as time-dependent changes of bone-related miRNAs in serum including associations to bone microstructure under anti-resorptive and osteo-anabolic therapy, respectively.

Given the large number of samples and data generated, the primary objective of the present study was to evaluate the expression of bone-related miRNAs in bone and serum under parathyroid hormone (teriparatide) or zoledronic acid (zoledronate) treatment in an OVX-model for postmenopausal osteoporosis after a full course of treatment. We hypothesize that: (i) ovariectomy induced bone-loss will result in specific changes of bone-related miRNAs compared to SHAM controls, (ii) that both zoledronic acid and teriparatide treatment might rescue these specific changes, and (iii) that the levels of these miRNA signatures in peripheral blood are correlated to bone tissue levels based on their release from bone cells.

## 2. MATERIALS & METHODS

This study was conducted at the Ludwig Boltzmann Institute for Traumatology, Vienna, Austria in cooperation with TAmiRNA GmbH (Vienna, Austria). The present study is part of the “TAMIBAT-Project” (**<u>T</u>**ime-dependent **<u>A</u>**nalysis of **<u>mi</u>**croRNAs and **<u>B</u>**one Microstructure under Consideration of **<u>A</u>**nti-Osteoporotic **<u>T</u>**reatment) which investigates the role of anti-osteoporotic treatment on bone-related miRNAs in bone tissue and serum.

### 2.1. Experiment Design

In total, 42 Sprague Dawley rats were either ovariectomized (OVX) or sham-operated at the age of 6 months. Eight weeks following OVX treatment, 32 animals were randomly assigned into three groups: (i) OVX + vehicle (VEH), n= 10; (ii) OVX + teriparatide (TPD), n=11; or (iii) OVX + zoledronic acid (ZOL), n=11. A fourth group was (iv) sham-operated (SHAM), n=10, and received placebo parallel to the OVX groups. Animals in group (i) received zoledronic acid (100 μg/kg/body weight s.c., single dose, Novartis Pharma AG). Animals in group (ii) received teriparatide s.c. periodically (weekly dose 210 μg/kg/body weight (= 42 μg/kg/body weight/per day over 5 days), Eli Lilly and Company). Group (iii) and (iv) acted as controls and received vehicle injections (0.9% saline with 20 mM NaH_2_PO_4_ in 0.9% NaCl, 3 mg/ml mannitol - dosed at a volume of 1 ml/kg body weight) s.c. at the same interval as group (ii).

### 2.2. Blood Collection

Blood was taken at the tail vein at week 0, 2, 8, 10, 12, 16 and 20. Blood sampling was performed in the early afternoon after a 6-hour fasting period to avoid influences of nutrition on bone metabolism. Blood serum was sampled in Eppendorf tubes from the lateral tail vein under short inhalation anesthesia (isoflurane 1 – 3%, oxygen). Clotting time for the serum samples was 30 – 45 minutes at room temperature. Samples were then centrifuged at 2500 × g for 15 minutes at 20°C. The same inhalation anesthesia protocol was used during in vivo micro-CT measurements. Study duration was 20 weeks for all subgroups, of which the treatment period accounted for 12 weeks. The complete study design is shown in Figure 1A.

**Figure 1.**
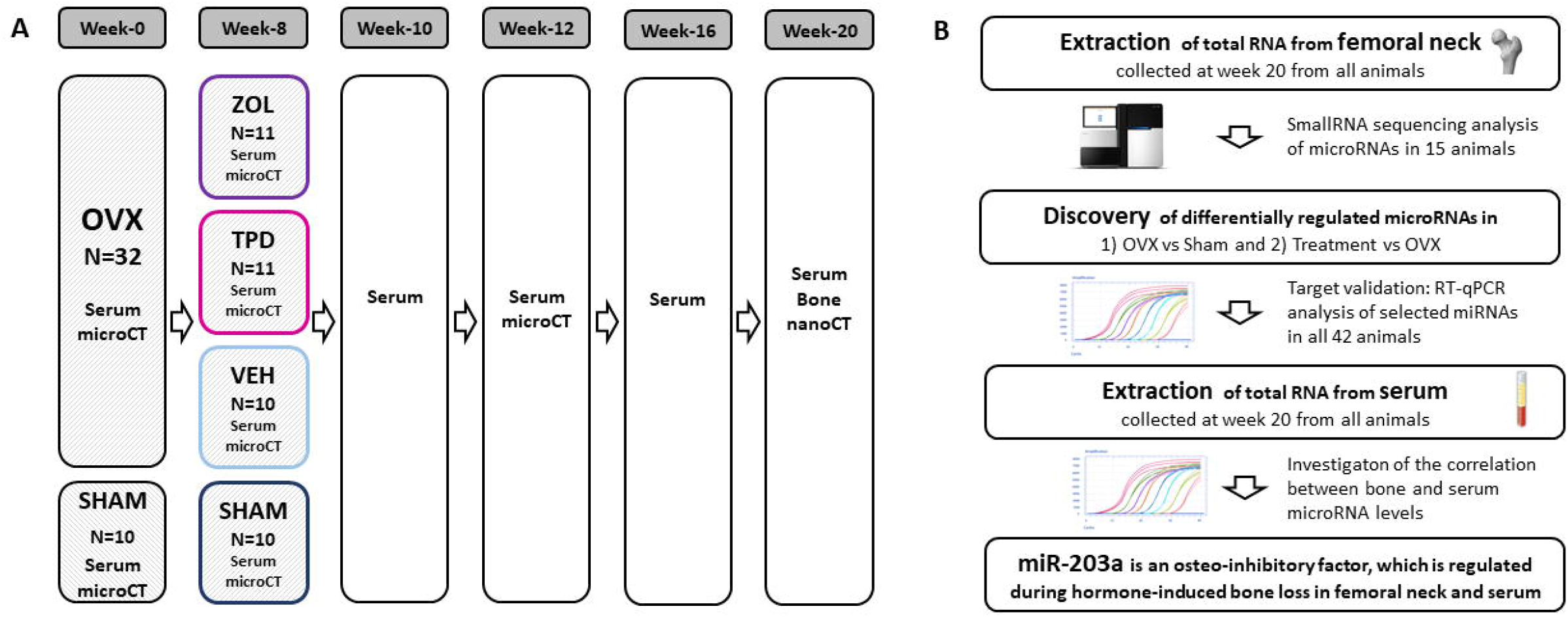
Overview of experiment design and microRNA analysis workflow of the TAMIBAT study. A) In total 42 Sprague Dawley rats at age 6-months underwent ovariectomy (OVX, n=32) or sham-surgery (SHAM, n=10). After 8 weeks, animals in OVX group were randomized to placebo treatment with vehicle solution (VEH, n=10), teriparatide treatment (TPD, n=11), or zoledronate treatment (ZOL, n=11). Micro-CT, nano-CT, serum and tissue analyses were performed as indicated. B) The microRNA analysis workflow in this study: initially total RNA was extracted from the femoral head of 42 animals. A subset of 15 animals (SHAM, n=6; VEH, n=3; TPD, n=3; ZOL, n=3) was used for small RNA sequencing. Differentially regulated miRNAs were confirmed by RT-qPCR and targeted analysis of microRNAs in matched serum samples was performed.

### 2.3. Animals

Thirty-two ovariectomized and 10 sham-operated 6-month-old female Sprague Dawley rats (ranging in weight from 270 g to 356 g) were sourced from Charles River Laboratories, Germany for this study. Animals were kept in makrolon type 4 cages with two animals per cage. They were fed ready-made feed (Rat Maintenance, Ssniff GmbH) and water ad libitum. Bedding was obtained from Abedd GmbH (Midi Chips). Gnawing blocks and tunnels were provided as cage enrichment. A sample size calculation was carried out based on primary variable bone density. Effect sizes from previous studies ^(23,24)^ were used in the calculation. A difference in the group average of 15% and a power of 90% was assumed, a group size of 10 was, therefore, adequate. Analogous studies have reported similar group sizes ^(23,24)^. All experimental protocols were approved in advance by the Municipal Government of Vienna in accordance with Austrian law and the Guide for the Care and Use of Laboratory Animals as defined by the National Institute of Health (revised 2011).

### 2.4. MicroRNA Analysis Serum

#### 2.4.1. RNA Extraction

Total RNA was extracted from 200 μl serum using the miRNeasy Mini Kit (Qiagen, Germany). Samples were thawed on ice and centrifuged at 12,000 x g for 5 minutes to remove any cellular debris. For each sample, 200 μL of serum were mixed with 1000 μL Qiazol and 1 μL of a mix of 3 synthetic spike-in controls (Exiqon, Denmark). After a 10-minute incubation at room temperature, 200 μL chloroform were added to the lysates followed by cooled centrifugation at 12,000 × g for 15 minutes at 4°C. Precisely 650 μL of the upper aqueous phase were mixed with 7 μL glycogen (50mg/mL) to enhance precipitation. Samples were transferred to a miRNeasy mini column where RNA was precipitated with 750 μL ethanol followed by automated washing with RPE and RWT buffer in a QiaCube liquid handling robot. Finally, total RNA was eluted in 30 μL nuclease free water and stored at −80°C to await further analysis.

#### 2.4.2. MicroRNA reverse-transcription quantitative PCR Analysis in serum RNA (RT-qPCR)

Starting from total RNA samples, cDNA was synthesized using the Universal cDNA Synthesis Kit II (Exiqon, Denmark). Reaction conditions were set in accordance to the manufacturer’s specifications. In total, 2 μL of total RNA were used per 10 μl reverse transcription (RT) reaction. To monitor RT efficiency and presence of impurities with inhibitory activity, a synthetic RNA spike-in (cel-miR-39-3p) was added to the RT reaction. PCR amplification was performed in a 96-well plate format in a Roche LC480 II instrument (Roche, Germany) using EXiLENT SYBR Green mastermix (Exiqon, Denmark) with the following settings: 95°C for 10 min, 45 cycles of 95°C for 10 s and 60°C for 60 s, followed by melting curve analysis. To calculate the cycle of quantification values (Cq-values), the second derivative method was used. Spike-in control values were used for monitoring data quality (Supporting Figure 1). Cq-values of endogenous miRNAs were normalized to the RNA spike-in controls by subtracting the individual miRNA Cq-value from the RNA Spike-In Cq, thus obtaining delta-Cq (dCq) values that were used for the analysis. Hemolysis was assessed in all samples using the ratio of miR-23a-3p versus miR-451a and applying a cut-off of a delta Cq > 7 indicates a high risk of hemolysis ^(25)^

### 2.5. MicroRNA/mRNA Analysis in Bone Tissue

#### 2.5.1. RNA Extraction

Femur bones were harvested from all rats. Any attached tissue was quickly removed and samples were immediately snap frozen in liquid nitrogen and stored at −80°C. For RNA extraction, only femoral heads of femur bones were used. For tissue homogenization, frozen bones were again put into liquid nitrogen and each bone was subsequently taken out and fixed in a jaw vice. Femoral heads were rapidly dissected at the femoral neck using a gigli wire saw, weighed and put back into liquid nitrogen. Femoral head samples were homogenized as described by Carter et al. ^(26)^. Each bone sample was added to a MPBio tube prechilled in liquid nitrogen containing Lysing Matrix M (MPBiomedicals, USA) followed by addition of 1000 μl Qiazol. Tubes containing the bone sample, Lysing Matrix M and 1000 μl Qiazol were put back in liquid nitrogen. Bone samples were then lysed by bead disruption in a MPBio Fastprep 24 instrument (MPBiomedicals, USA). After lysis, the tubes were centrifuged at 8600 × g for 30 seconds and the resulting supernatant was used for total RNA extraction using the miRNeasy Mini kit (Qiagen, Germany) as per the manufacturer’s recommendation.

#### 2.5.2. Small RNA Sequencing

Equal amounts of total RNA (100 ng) were used for small RNA library preparation using the CleanTag small RNA library preparation kit (TriLink Biotechnologies). Adapter-ligated libraries were reverse transcribed and amplified (19 cycles) using barcoded Illumina reverse primers in combination with the Illumina forward primer according to the instructions of the manufacturer. In total 15 small RNA sequencing libraries were prepared from bone tissue RNA extracts and used for equimolar pooling based on DNA-1000 high-sensitivity bioanalyzer results (Agilent Technologies). The library pool was sequenced on a NextSeq 500 sequencing instrument (50 bp cycles) following the manufacturer’s instructions. Raw data was de-multiplexed and FASTQ files for each sample was generated using the bcl2fastq software (Illumina inc.). FASTQ data were checked using the FastQC tool (http://www.bioinformatics.babraham.ac.uk/projects/fastqc/). Sequencing reads were adapter trimmed and filtered for low-quality reads (Q < 30). MicroRNA annotation was performed on the basis of sequential alignments against the rat genome reference and miRBase release 21. Read counts were normalized to the total number of reads detected per sample to obtain the “tags per million” (TPM) for each miRNA and sample. Exploratory data analysis was performed using Clustvis (https://biit.cs.ut.ee/clustvis/) ^(27)^.

#### 2.5.3. MessengerRNA (mRNA) reverse-transcription quantitative PCR Analysis (RT-qPCR)

For mRNA quantification the TATAA SYBR Grandmaster mix kit (TATAA Biocenter, Sweden) was used. 25 ng of bone tissue total RNA extracts were used for reverse transcription and all steps were carried out according to recommendations by the manufacturer. PCR amplification was performed in a 96 well format in a Roche LC480 II instrument (Roche, Germany) with the following settings: 95 °C for 30 seconds followed by 45 cycles of 95 °C for 5 seconds, 63 °C for 15 seconds and 72 °C for 10 seconds and subsequent melting curve analysis. To calculate the cycle of quantification values (Cq-values), the second derivative method was used.

**Table 1:**
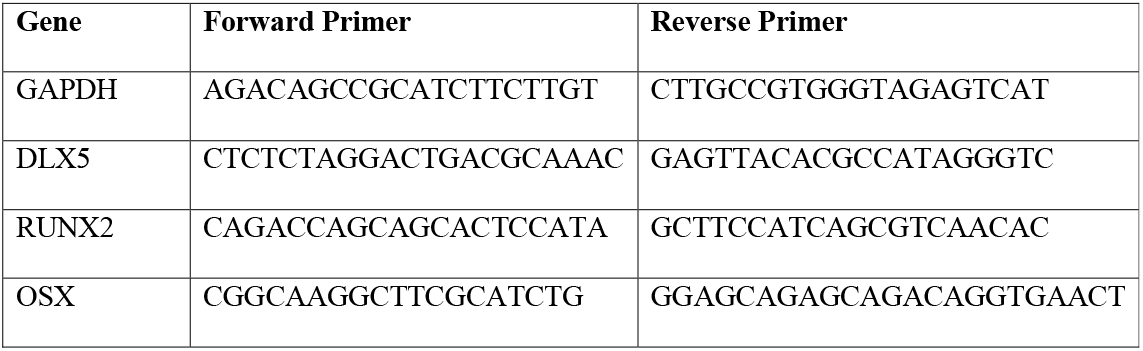
Primer sequences used for mRNA reverse-transcription quantitative PCR analysis.

#### 2.5.4. MicroRNA reverse-transcription quantitative PCR Analysis in bone RNA (RT-qPCR)

Bone microRNA analysis was performed similarly to serum microRNA analysis with the following modifications: precisely 10 ng of total RNA were used for reverse transcription. Variance in reverse transcription was analyzed using spike-in controls (cel-miR-39). To account for variance in reverse transcription, qPCR data were normalized to cel-miR-39 (delta-Cq method).

### 2.6. Bone Microstructure Assessment

Micro-CT-scans were performed at the vertebral body L4 and the tibia in all animals. In-vivo μCT scans were performed using a SCANCO vivaCT 75 (SCANCO Medical AG, Brüttisellen, Switzerland) at week 0 (baseline), 8 and 12. Scans were performed at 70 kVp, 114 μA, 500 projections/180° with an integration time of 300 ms and reconstructed to an isotropic resolution of 45 μm.

The bone microstructure at study end (week 20) was evaluated using a μCT imaging system (nano-CT, μCT 50, SCANCO Medical AG, Brüttisellen, Switzerland). The tube was operated at 90 kVp, 200 μA, 1000 projections/180° with an integration time of 500 ms. The scans were performed at an isotropic nominal resolution of 10 μm. The long axis of the biopsy specimen was oriented along the rotation axis of the scanner.

Image segmentation was performed using Definiens Developer XD 2.1.1 (Definiens AG, Munich, Germany). The tibia scans were downsized to 12.5% and maintained at 375 mg HA/cm^3^. Void volume inside and outside of the bone was separated by coating the bone tissue 2 voxel (160 μm) thick with a temporary class. Void volume in contact with the image border was classified as exterior void and void enclosed by bone was classified as interior void. The temporary coating was removed by growing the void classes into the temporary class. The fibula was excluded from further measurements. As the cross-section area of the fibula was categorically smaller than the cross-section area of the rest of the tibia, it could be automatically distinguished and excluded. Cortical and trabecular bone was distinguished using a series of surface tension constrained region growing steps. The exterior void was coated as cortical bone into bone constrained by surface tension. This resulted in a separation of the dense outer portion of the bone (the cortex) from the less dense trabecular bone.

The vertebra was imported into Fiji and the region of interest was drawn to include the trabecular space in the vertebral body by using the polygon tool with interpolation using the ROI manager. A mask was created from the ROI and imported into Definiens Developer XD 2.1.1 together with the original scan. The region of interest was reduced to 5 pixels (50 μm) in all directions to ensure that no parts of the cortical bone were included and segmented using a threshold of 700 mg HA/cm^3^. The segmentation was exported as image stacks using black for enclosed void, white for bone and indicating outside of the region of interest. The segmented images were then imported in Fiji and measured using the BoneJ plugin.

### 2.7. Statistical Analysis

Statistical differences in bone microstructure, bone microRNA/mRNA transcript levels, and serum microRNA transcript levels between SHAM and the three OVX groups were conducted using GraphPad Prims v5.03. Statistical significance of group differences was assessed using 1-way-ANOVA in conjunction with Tukey post hoc tests for pair-wise comparisons. Scatterplots present the distribution of values as well as mean +/- standard deviation. Statistical analysis of small RNA sequencing data based on TPM values was performed using the methods available under the EdgeR package in R/Bioconductor. P-value obtained from EdgeR were adjusted for multiple testing using Benjamini-Hochbergs false discovery rates (FDR) ^(28)^.

## 3. RESULTS

### 3.1. Study aim and experiment design

The aim of the TAMIBAT study is the systematic investigation of microRNA levels in rodent bone tissue and cell-free blood (serum) during the progression of hormone-induced bone loss and in response to anabolic and anti-resorptive rescue of bone loss, respectively. Figure 1 summarizes the experiment’s design: 42 six-month-old Sprague Dawley rats were included into the study and randomized to OVX or SHAM treatment. Eight weeks after surgery, OVX animals were randomized to treatment with vehicle solution (VEH), teriparatide (TPD), or zoledronate (ZOL). Treatments as well as multiple blood withdrawals and in vivo micro-CT analysis were performed within a time frame of 12 weeks, ending with sacrifice of all animals and collection of blood and bone tissue for gene expression and ex vivo nano-CT analysis (Fig. 1A). Initially, we were interested in the effects of bone loss as well as therapy on bone tissue microRNA levels. Therefore, we applied next-generation sequencing (NGS) for the discovery of changes in small RNAs, specifically miRNAs, on tissue level and RT-qPCR to confirm these findings in the complete sample set (Fig. 1B).

### 3.2. Bone Microarchitecture

We observed a significant reduction in trabecular bone at week 20 in ovariectomized and untreated animals (VEH) at sites of tibial and vertebral bone (Fig 2). Anabolic treatment (TPD) resulted in a complete rescue of bone loss after 20 weeks, while anti-resorptive treatment (ZOL) resulted in only a partial rescue of BV/TV. These effects were similarly observed at both the tibia and vertebral bone sites.

**Figure 2.**
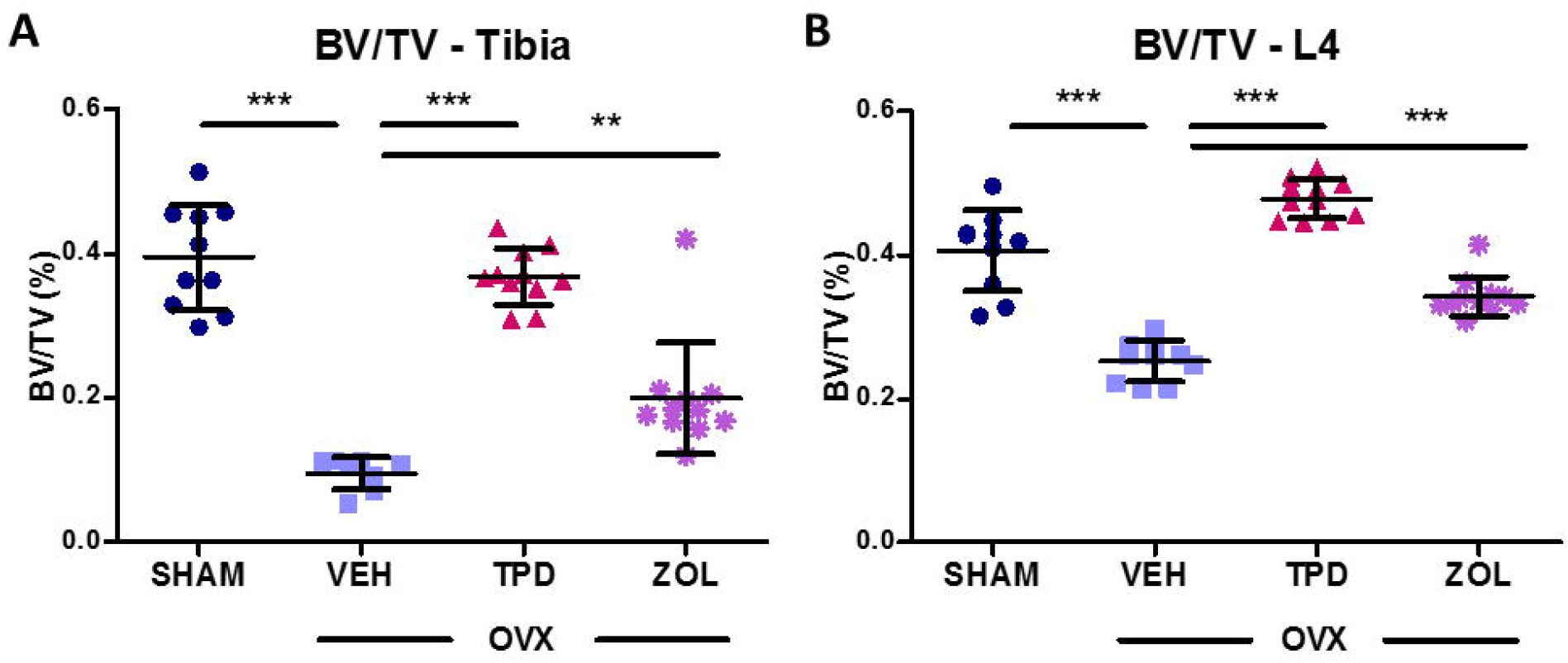
Bone microstructure analysis by micro-CT and nano-CT. A/B) Ex vivo nano-CT analysis of the effects on tibial and vertebral bone measured as BV/TV (SHAM, n =10; VEH n=10, TPD, n=11; ZOL n=11). Scatterplots show mean +/- SD. Testing was performed using 1-way ANOVA and Tukey post hoc test for pairwise comparisons. * p<0.05, ** p<0.01, *** p<0.001.

### 3.3. MicroRNA expression in femoral head in response to anti-osteoporosis therapy

To investigate changes in bone microRNA expression as a consequence of ovariectomy (OVX) and in response to treatment, femoral head samples were harvested at week 20 and used for RNA extraction. Initially, NGS was used for a genome-wide screen of miRNA levels in a subset of 15 out of 42 samples (6x SHAM, 3x VEH, 3x TPD, 3x ZOL). Samples were selected based on nano-CT results. NGS detected 426 distinct miRNAs in femoral head tissue, which yielded at least one tag per million (TPM) in at least one out of the 15 samples. Out of the 426 detected miRNAs, 273 gave one or more TPMs in each of the 15 samples and were subsequently used for statistical analysis. Using TPMs from the most variant microRNAs (top 30 sorted by CV%) a heatmap was generated (Supporting Figure 2), which showed distinct microRNA signatures in vehicle treated OVX animals that was clearly different to that of SHAM, TPD and ZOL animals. In contrast, miRNA expression in bone of SHAM and treated animals was observed to be more similar. Next, we performed statistical analysis of the group differences using EdgeR. P-values were adjusted for multiple testing and an adjusted p-value cut-off of 0.1 (FDR = 10%) was applied to identify differentially expressed miRNAs. Comparing vehicle-treated OVX animals (VEH) to SHAM controls we identified 13 significantly up-regulated miRNAs in VEH compared to SHAM control, while no miRNA was down-regulated (Fig. 3A). Anabolic treatment using teriparatide had profound effects on bone miRNA expression compared to vehicle-treated controls (VEH) and resulted in a significant regulation of 46 miRNAs (21 up, 25 down) (Fig. 3B), while 10 miRNAs (1 up, 9 down) were found to be regulated between ZOL and VEH (Figure 3C). Seven miRNAs (miR-203a-3p, miR-17-5p, miR-335-5p, miR-132-3p, miR-34a-5p, miR-19a-3p, miR-181b-3p) which showed significant differential expression in any of these comparisons were successfully replicated by RT-qPCR to confirm the validity of NGS results (Supporting Figure 3).

**Figure 3.**
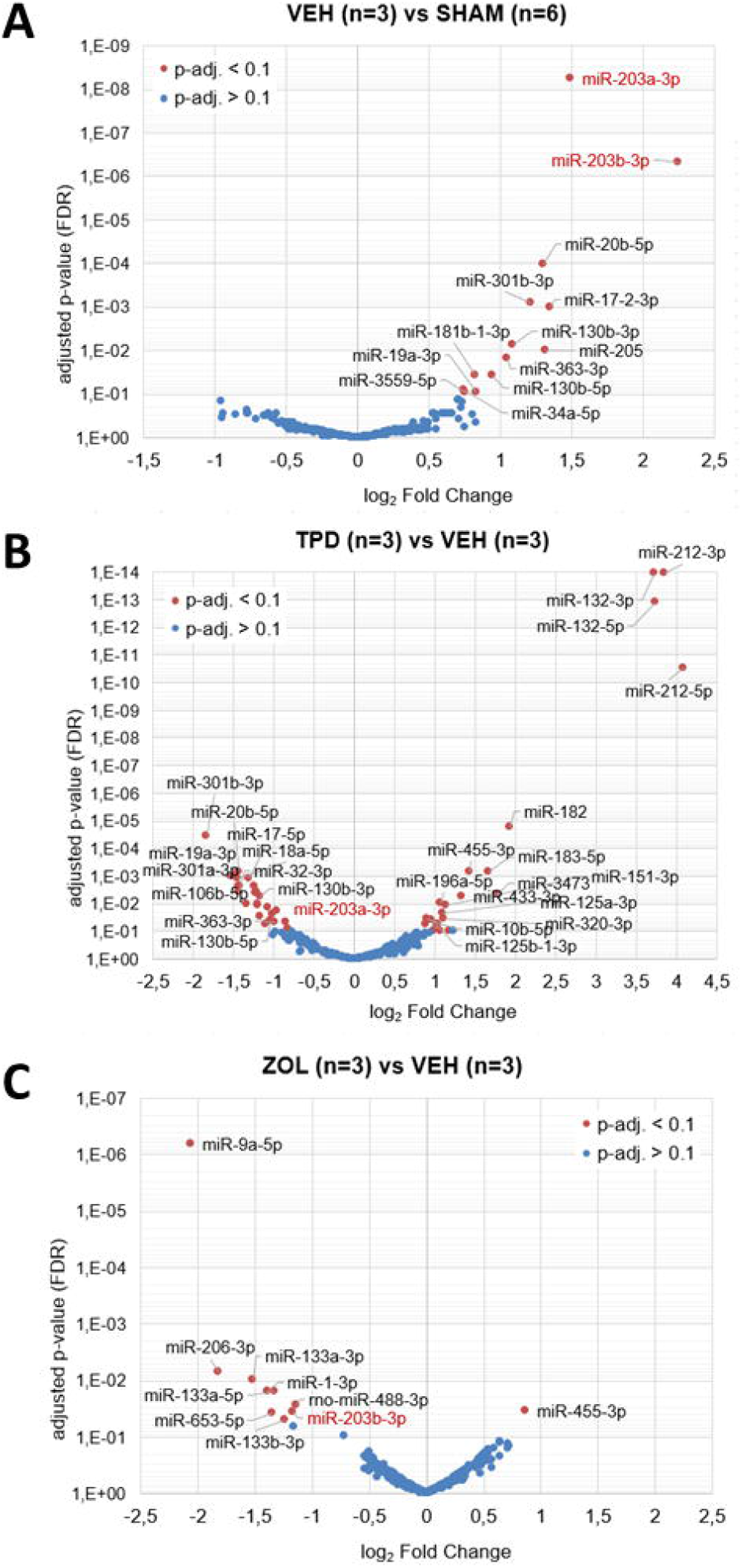
NGS-discovery of microRNA changes in femoral head bone tissue of ovariectomized animals as well as animals undergoing anti-osteoporotic treatment. A-C) Volcano plots depict the log2 transformed fold change and adjusted p-values for three contrasts *(SHAM, n=6; VEH, n=3; TPD, n=3; ZOL, n=3)*. microRNA effects with Benjamini-Hochberg adjusted p-values<0.1 (FDR < 10%) are highlighted as separate group. miR-203a/b is highlighted in red.

### 3.4. miR-203a expression is up-regulated in bone of ovariectomized animals and rescued by treatment

In the NGS screen miR-203a/b-3p showed the highest up-regulation in bone of OVX animals receiving vehicle (VEH) treatment compared to SHAM controls (log_2_FC = 1.5, FDR < 0.001, see Fig. 3A). Conversely, anabolic treatment with teriparatide led to a significant down-regulation of miR-203a in femoral head compared to vehicle-treated animals (log_2_FC = −0.85, FDR = 0.045, see Fig. 3B). Similarly, zoledronate (ZOL) treated femoral head bone tissue exhibited lower miR-203b expression (log_2_FC = −1.17, FDR = 0.04). Accordingly, we selected miR-203a for further investigation.

Levels of miR-203a were first analyzed in the entire cohort of 42 animals by RT-qPCR to confirm NGS results and increase sample size. Up-regulation of miR-203a in VEH versus SHAM group and rescue due to teriparatide treatment were confirmed (p<0.05, Fig. 4A). Although treatment with zoledronic acid did not result in a rescue of miR-203a levels, levels were lower by trend compared to VEH animals (Fig. 4A). RT-qPCR was used to quantify mRNA levels of two transcription factors that are regulated by miR-203a, namely Dlx5 and Runx2, as well as Osterix (Osx), which acts down-stream of Runx2. The regulation of Dlx5 was inverse to miR-203a, indicated by down-regulation in the OVX group with vehicle treatment (VEH) and up-regulation upon teriparatide treatment (TPD) (p < 0.05, Fig. 4B). Likewise, Runx2 was observed to be down-regulated in bone of untreated OVX animals (VEH) compared to SHAM (Fig 4C). However, treatment with teriparatide did not result in an induction of Runx2 mRNA levels. Osterix mRNA levels showed the same trend as Runx2 (r=0.84, p<0.001) but in contrast to Runx2, the down-regulation in VEH vs SHAM did not reach significance.

**Figure 4.**
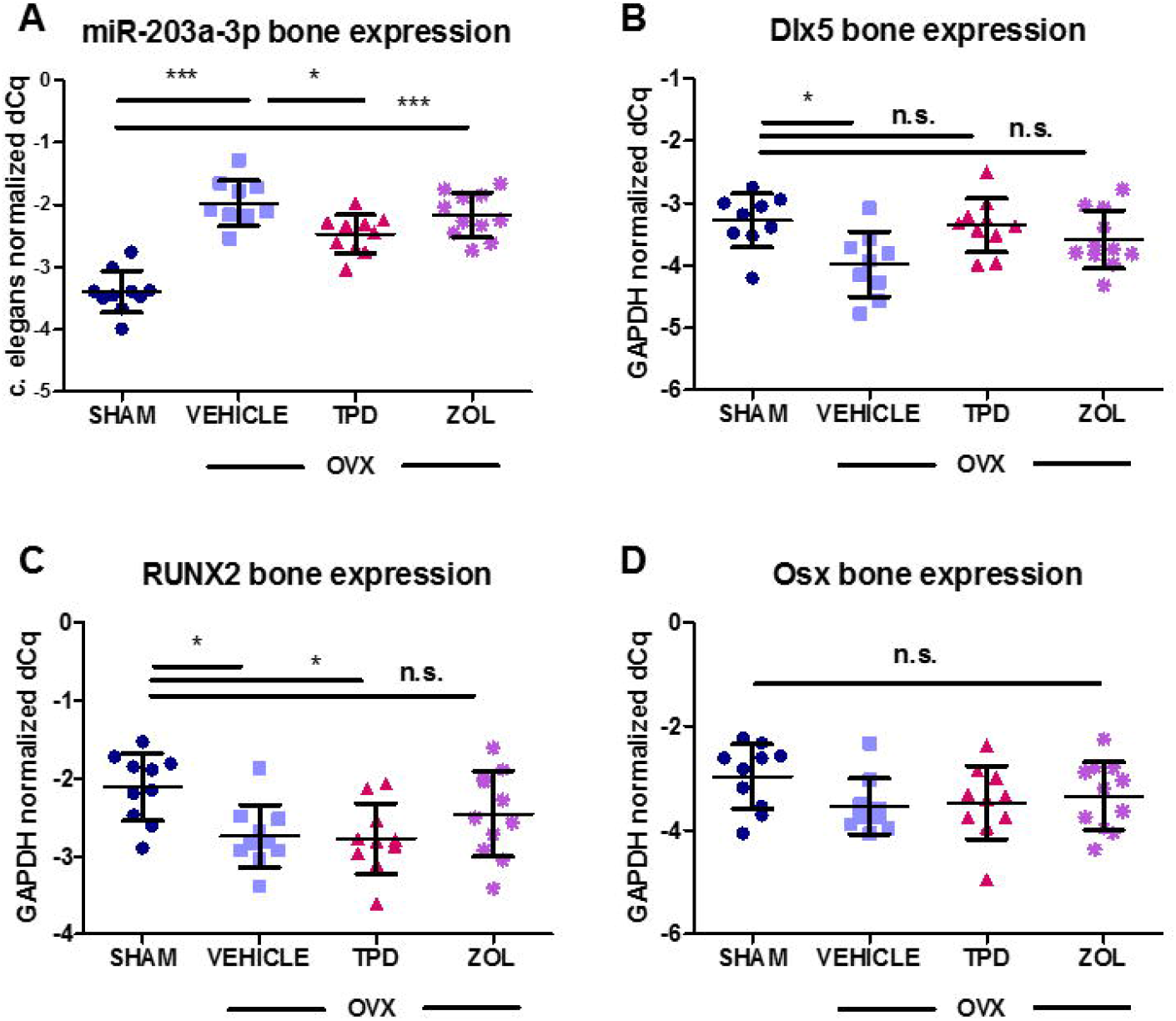
Regulation of miR-203a-3p and confirmed target genes in femoral head bone tissue of the entire TAMIBAT cohort. (SHAM, n=10; VEH, n=10; TPD, n=10; ZOL, n=11). A) RT-qPCR quantification of miR-203a. B) RT-qPCR quantification of Dlx5. C) RT-qPCR quantification of Runx2. D) RT-qPCR quantification of Osx. Scatterplots show mean +/- SD. Testing was performed using 1-way ANOVA and Tukey post hoc test for pairwise comparisons. * p<0.05, ** p<0.01, *** p<0.001.

### 3.5. Serum levels of miR-203a are significantly correlated to expression in bone

In order to monitor circulating levels of miR-203a after 12 weeks of treatment, total RNA was extracted from serum samples of all 42 animals included in the TAMIBAT study at week 20. We observed a significant correlation (r=0.41, p<0.05) between miR-203a-3p levels in bone and matched serum samples (Fig. 5).

**Figure 5.**
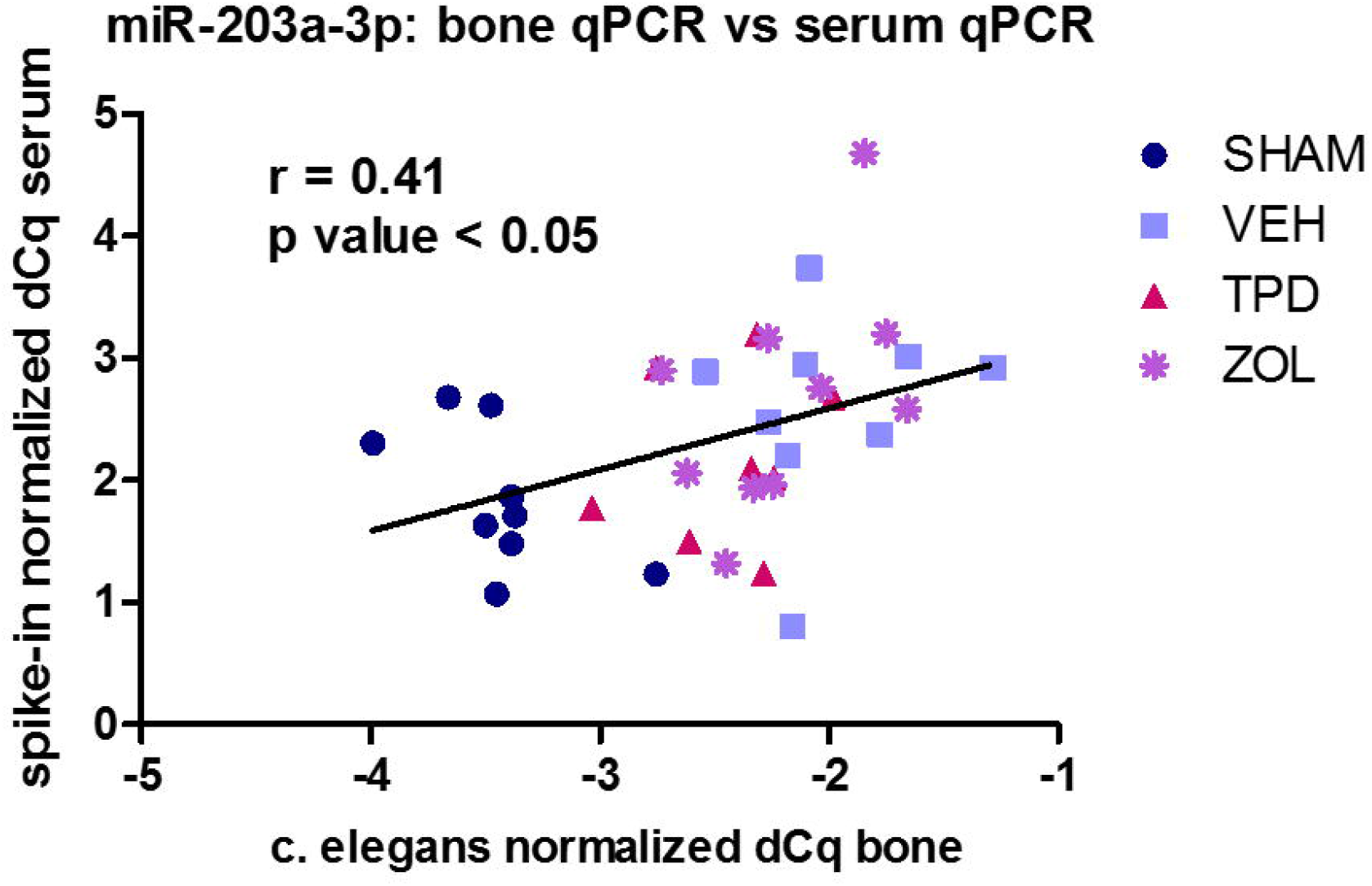
Correlation between bone (x-axis) and serum (y-axis) levels of miR-203a-3p based on n=36 (SHAM, n=9; VEH, n=9; TPD, n=8; ZOL, n=10) animals in the TAMIBAT study.

## 4. DISCUSSION

In this classical animal model for postmenopausal osteoporosis, the expression of bone-related miRNAs was investigated under teriparatide and zoledronic acid treatment, respectively. Although hormone replacement therapy (HRT) shows advantageous effects on bone metabolism, it is not considered first line therapy for the treatment of osteoporosis. Therefore, the aim of the present study was to investigate the effects of an established antiresorptive or an osteo-anabolic agent, as the mode of action as well as the effects on bone mineral density and fracture risk reduction are different ^(29)^.

Total RNA was extracted from femoral head tissue, as this bone can be collected quickly and reproducibly, thus enabling the ability to obtain high quality RNA from bone cells and to a lesser extent bone marrow cells. Circulating miRNAs were analyzed using cell-free blood (serum) of ovariectomized rats and SHAM controls. Ovariectomy without therapy, reflecting postmenopausal osteoporosis, as well as anti-resorptive and osteo-anabolic treatment resulted in distinct miRNA changes in bone tissue. In total, 11 miRNAs were up-regulated in untreated OVX animals and down-regulated under a teriparatide or zoledronate treatment regimen. These miRNAs seem be mechanistically relevant regarding postmenopausal bone loss and treatment response.

miR-203a showed the most significant up-regulation in bone of ovariectomized untreated rats compared to SHAM, which was rescued by teriparatide treatment. miR-203a was previously described to be an important regulator of bone formation, as it is involved in the differentiation of mesenchymal stem cells. For example, down-regulation of miR-203a promotes osteoblastic differentiation by up-regulation of osteogenic effectors including PPAP2B, NEGR1 and PDGFRA ^(30)^. In addition, miR-203a acts as a potent inhibitor of Dlx5, which regulates the transcription factors Osx and Runx2. A direct interaction between miR-203a and Runx2 has been proposed in a breast cancer model ^(31)^. Consequently, osteoblast differentiation is slowed by miR-203a ^(32)^. In line with these previous observations, elevated expression of miR-203a was found in the present study in femoral head tissue and serum of vehicle-treated OVX rats, representing the untreated postmenopausal osteoporotic phenotype, suggesting that miR-203a might contribute to delayed osteogenic differentiation and hence development of bone loss. Contrarily, inhibition of miR-203a in vitro resulted in the stimulation of matrix mineralization in the study of Laxman et al. ^(32)^. Laxman et al. have shown that the transcription of miR-203a itself seems to be negatively controlled by BMP2, while it can be induced through treatment with dexamethasone (Fig. 6) ^(32).^ Similar to the previously described up-regulation of miR-203a in cell-free blood of untreated OVX rats, miR-203a was also found to be up-regulated in serum of diabetic and non-diabetic postmenopausal women with osteoporosis and a history of low-traumatic fractures ^(17)^. Taken together, these findings suggest an important role for miR-203a in the regulation of bone formation and potentially during the onset and development of postmenopausal osteoporosis. In bone samples of untreated ovariectomized rats we observed that both Dlx5 and Runx2 were down-regulated in comparison to SHAM animals. However, low Runx2 levels were also found in the teriparatide (TPD) treated group. These findings are somewhat surprising as TPD is an anabolic agent and bone microstructure and density increased significantly following 12-weeks of treatment, suggesting a sufficient treatment response. The increased transcription of Runx2 during bone formation is believed to be oscillating rather than constant, indicating that the time point of analysis may be essential to detect up-regulation. In addition, mRNA and protein levels are not necessarily correlated, especially due to the action of post-transcriptional regulators such as miRNAs. For example, a study on TPD-treated osteoblastic MC3T3-E1 cells observed an unexpected down-regulation of Runx2 as well. In this study, femoral bone tissue was taken from rats 48 hours after the last treatment with TPD, which might be responsible for the observed effect. It has to be taken into account, that the mechanisms of osteo-anabolic drugs are complex. In addition to Runx2 other mechanisms including the Wnt/β-catenin pathway are responsible for osteoblast differentiation under TPD-treatment ^(33)^. miR-203a might be an important regulator of osteoblast differentiation. However, it does not reflect the whole body microenvironment.

**Figure 6.**
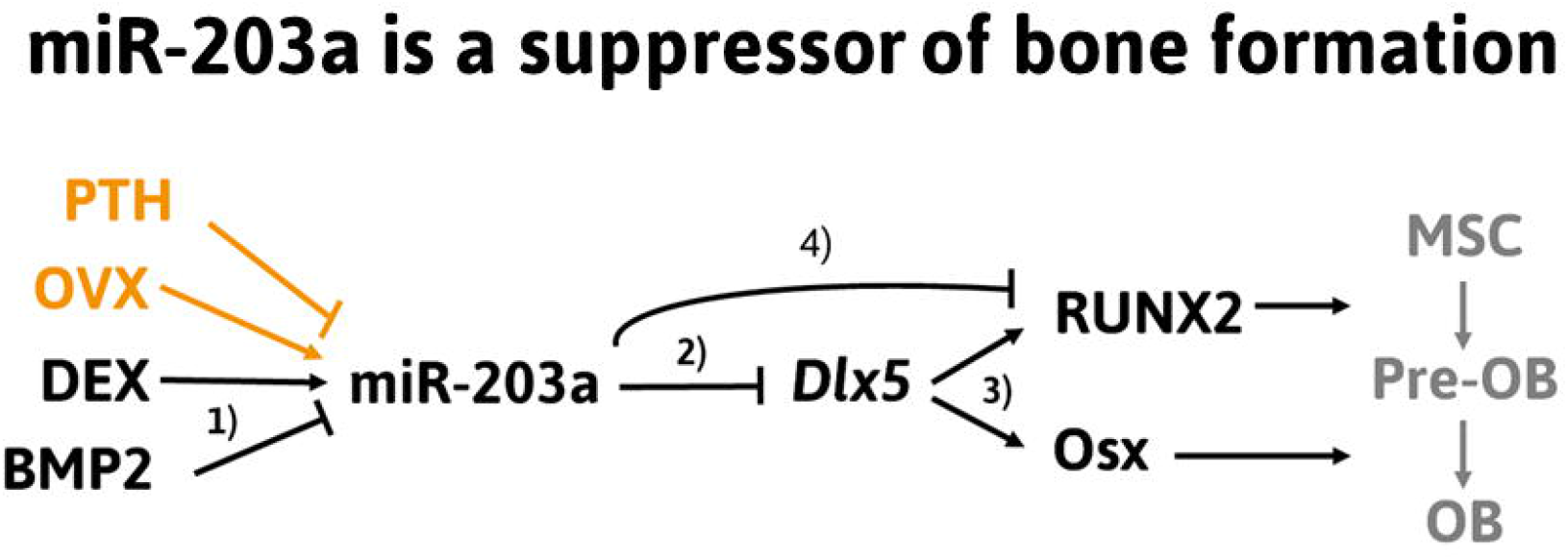
miR-203a is a suppressor of bone formation. PTH, parathyroid hormone; OVX, ovariectomy, DEX, dexamethasone; BMP2, bone morphogenic protein 2. Orange colors indicate the effects that were discovered in this study. References: 1) Laxman N. et al., Bone 2016; ^(39)^ 2) Laxman N. et al. Genes (Basel) 2016; ^(32)^ 3) Lee M. et al. Biochem Biophys Res Commun.2003; ^(40)^ 4) Taipaleenmäki H. Cancer Res. 2015 ^(31)^.

Another key observation made by this study is that miR-203a levels in bone tissue and serum are significantly regulated. This finding confirms that changes in miRNA levels in bone tissue can be detected using liquid biopsies. A similar concomitant up- or down-regulation of miRNAs in bone tissue, serum as well as in osteoblasts and osteoclasts was also reported by other study groups ^(34,35)^ and encourages the potential clinical utility of miRNAs as bone biomarkers.

Apart from miR-203a, miR-20a-5p levels were also up-regulated following ovariectomy and down-regulated due to TPP therapy. miR-20a-5p was recently reported to be one of the most abundant miRNAs in bone tissue from postmenopausal women ^(36)^. Similar to miR-203a, miR-20a-5p also regulates the activity of transcription factors of osteoblast differentiation such as Runx2, BMP2 and PPARγ ^(36,37)^ and thereby osteoblast differentiation and bone formation.

Currently no other data on miRNAs in models of bisphosphonate treated postmenopausal osteoporosis are available. Teriparatide and denosumab therapy changed circulating miRNAs in postmenopausal osteoporosis in the study of Anastasilakis et al. ^(22)^. In this study, miRNA-133 was down-regulated under teriparatide, but not denosumab therapy, suggesting an inhibitory role of miRNA-133 on the RUNX-2 gene and therefore osteoblastogenesis ^(22)^. In the present study, miRNA-133 was down-regulated in animals receiving zoledronic acid, suggesting a positive effect of bisphosphonates on RUNX2 and thus bone formation. Significantly different expressions of miRNAs, including bone-related miR-29b-3p, were found between estrogenic hormone replacement therapy users and their treatment-naïve twins ^(38)^. If estrogen replacement can rescue or reverse the effects of estrogen deficiency on miRNAs is an important issue and should be raised in further research projects.

## 5. CONCLUSION

By taking advantage of a standardized animal model of postmenopausal osteoporosis, profound effects of hormone-induced bone loss as well as osteo-anabolic and anti-resorptive treatment on miRNA expression in bone, specifically femoral head, were revealed. Therapeutic intervention, especially osteo-anabolic therapy, resulted in a restoration of miRNA expression similar to sham-operated animals. Among the affected miRNAs, the induction of miR-203a in bone of ovariectomized rats was most significant and, notably, also detectable in cell-free blood. The regulation of miR-203a also affected mRNA levels of its direct targets, the transcription factors Dlx5 and Runx2, in femoral head tissues.

## 6. ACKNOWLEDGEMENT

The authors acknowledge T.A. Vacca from Linz/Austria for proofreading.

## 7. CONTRIBUTION

Individual contributions include the following: R.K., M.H., P.P., J.G. and H.R. for the study concept and design; M.W., E.G. and S.S. for the data collection; P.H. and R.K. for μCT imaging; M.H., M.W., S.S. and E.G. for miRNA analysis; G.L. and J.F. for the animal care; R.K., M.H. and M.W. for the data analysis; R.K., M.H., P.P., H.R. and J.G. for the data interpretation; R.K., M.H. and M.W. for drafting the manuscript and literature research; J.G., H.R. and P.P. for the extensive revision of the manuscript.

---

**Supporting Figure 1. Quality Control of circulating microRNA analysis in rodent serum samples**. Raw Cq-values obtained for RNA spike-in control (UniSp5) and cDNA spike-in control (cel-miR-39) are plotted for all samples included in the data analysis (SHAM, n=9; VEH, n=9; TPD, n=9; ZOL, n=10).

**Supporting Figure 2. Heatmap**. Normalized NGS read counts (TPM-values) for the top 30 microRNAs (ranked according to their coefficient of variation) were used to draw a heatmap. Rows represent microRNAs, columns represent samples. Pearson correlation and complete linkage were used for clustering of samples.

**Supporting Figure 3. Confirmation of NGS data by RT-qPCR**. In total 15 animal bone tissue samples (SHAM, n=6; VEH, n=3; TPD, n=3; ZOL, n=3) were used for NGS analysis. For those samples, the log_2_-transformed TPM values from the sequencing experiment were plotted against the normalized delta-Cq values obtained from RT-qPCR analysis.

