## Supplementary figures and images for "MicroRNA levels in bone and blood change during bisphosphonate and teriparatide therapy in an animal model of postmenopausal osteoporosis"

### Supporting-Figure-S1

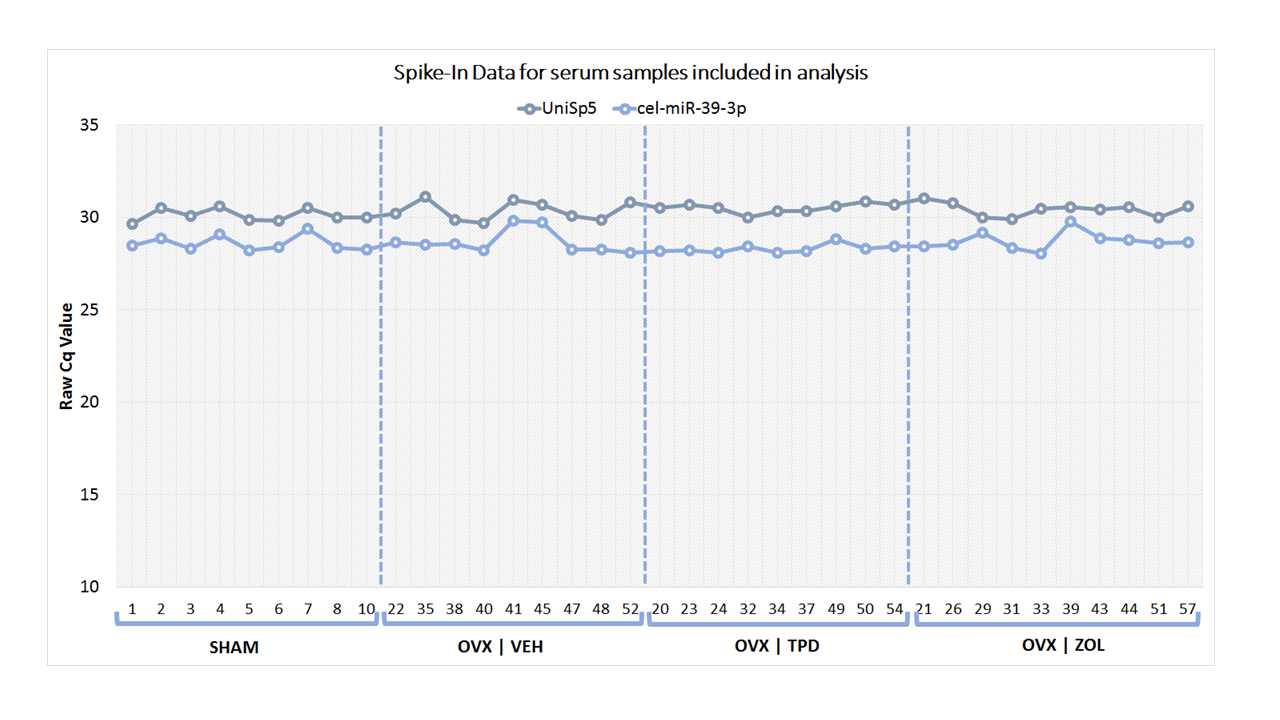

### Supporting-Figure-S2

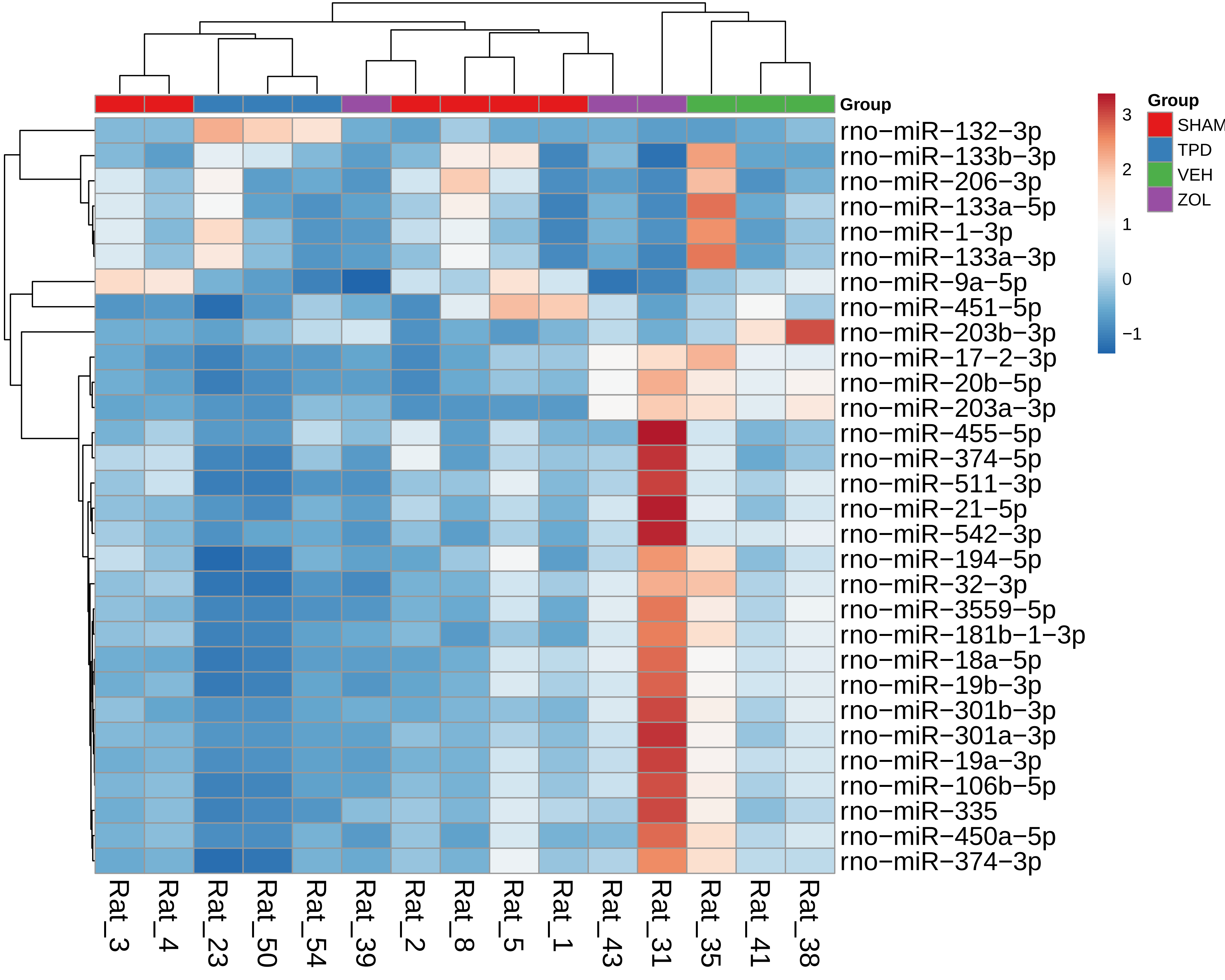

### Supporting-Figure-S3.1

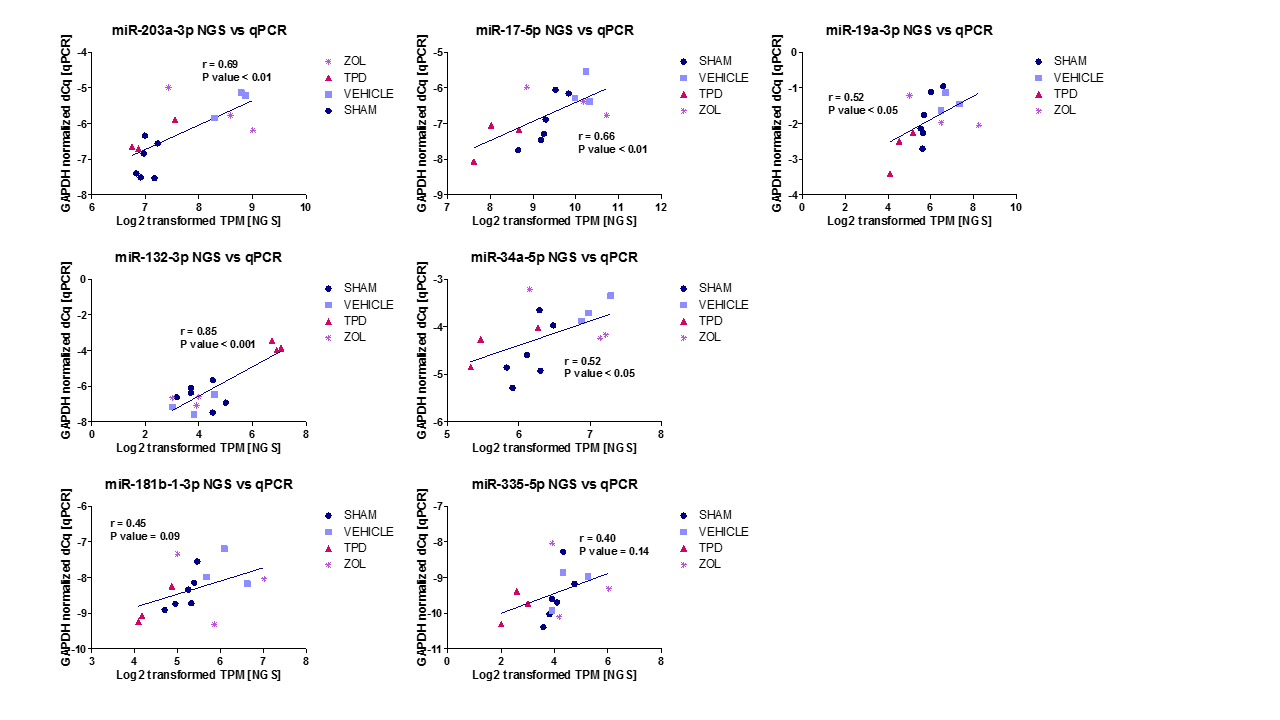

### Supporting-Figure-S3.2

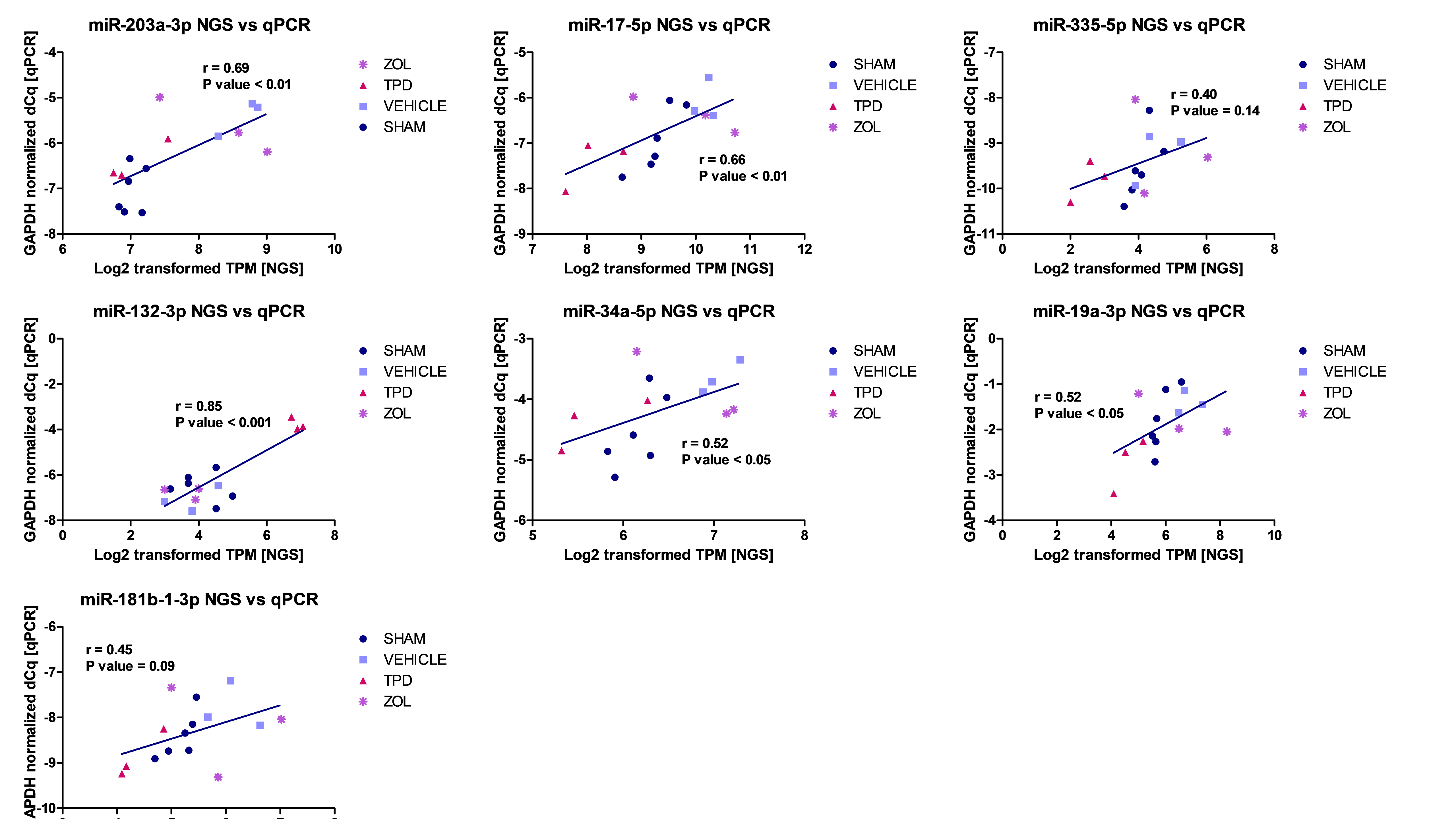
